# Pancreatic progenitor cells respond to laminar shear stress with transcriptomic changes, accompanied by reduced YAP1 activity

**DOI:** 10.64898/2026.09.29.754874

**Authors:** Xinren Yu, Ashwin Jiju, Rubitha Souresh, Maryam Hasani, Leonie Gäbler, Allison C. Lewis, Anne Grapin-Botton, Adele Doyle

## Abstract

The human pancreas secretes digestive fluid that flows through an extensive ductal network before emptying into the duodenum. However, little is known about how hydrodynamic forces influence pancreatic function and development. We hypothesized that pancreatic cells respond to physiological levels of fluid shear stress. In this study, we developed a microfluidic system featuring reversible sealing between the device layer and the cell culture substrate, enabling (1) conventional open-well cell culture prior to fluid shear stress application and (2) selective patterning of proteins and cells for migration studies. Channel dimensions were chosen to support 24-hour perfusion using standard syringe pumps. Using this platform, we investigated effects of physiological shear stress levels representing basal and activated pancreatic function (2, 8 and 33 mPa) on human iPSC-derived pancreatic progenitor cells. Pancreatic progenitor cells exhibit graded sensitivity to shear stress at the transcriptomic level. Specifically, shear stress upregulated expression of cell cycle-related gene sets; endocrine-associated genes including *NEUROG3*; and Notch receptor ligands (*DLL1*, *DLL3*) in a magnitude-dependent manner. The exposure to unidirectional fluid flow also resulted in a reduction in YAP1 protein content, decreased *WWTR1* gene expression, and downregulation of YAP/TAZ target genes. Furthermore, barrier-based migration assays under shear stress revealed that cell patches expanded their boundaries both along and against the flow, with greater distance in the direction of flow. Our system and findings may help lay the foundation for understanding biophysical regulation of developmental processes in a physiologically important, yet hard-to-access, epithelial model system.

## 1. Introduction

Fluid flow occurs throughout the human body and plays an important role in organ function, from maintaining circulation in blood vessels to supporting transport processes in the digestive system. Along the digestive tract, the pancreas secretes over 1 liter of fluid daily.^1^ Pancreatic juice is produced by the exocrine compartment, including acinar cells that release digestive enzymes and ductal cells that produce bicarbonate and the majority of the fluid, and this juice is delivered into the duodenum.^2,3^ The pancreas also contains endocrine cells that secrete hormones into the blood vessels to regulate glucose homeostasis. Although flow is a physical cue intrinsic to normal pancreas function, little is known about how flow itself may affect pancreatic ductal cells.

In addition to the importance of fluid transport for the adult pancreas function, it has been postulated that fluid flow promotes ductal remodeling from a network to a tree during pancreas development. Fluid flow also actively contributes to embryonic development in multiple organs, including left-right asymmetry establishment,^4^ heart valve formation,^5^ and vascular remodeling.^6,7^ Developing blood vessels undergo regression to form the mature vasculature; likewise, branch pruning has been observed in the developing pancreatic ductal network, as it transitions from a plexus structure with redundant ducts at E14.5 to a tree-like structure with hierarchy at E18.5 in mouse models.^8^ Within lumenal space, pancreatic secretion has been detected as early as E12.5 in mouse explants.^8^ However, whether fluid flow actively contributes to pancreatic development remains unclear.

As fluid flows past the ductal linings, cells experience shear stress. While the role of hydrodynamics is best established in vascular biology, recently shear stress effects on other epithelial cells have attracted considerable attention as well,^9–13^ including in the kidney,^14^ lungs,^14,15^ brain ventricles,^16^ or epididymis.^17^ Previous studies have applied μPa-level shear stress to mimic interstitial flow for reaggregated pancreatic islet clusters^18^ or Pa-level shear stress to study the role of Piezo1-mediated mechanosensitive calcium signaling in mature acini.^19^ However, in these two studies, the flow applied differed from the physiological context as islet flows occur primarily in fenestrated islet microvessels and in acini the shear was applied on the basal side rather than apical, where the exocrine fluid is secreted. Neither study was intended to recapitulate physiological fluid shear stress exerted on the apical surface within the pancreatic ductal lumen, where exocrine fluid is secreted and flows. Moreover, nothing is known about the cellular responsiveness to shear stress in the developing pancreas. The mechanobiology research in the last two decades has revealed that biophysical cues such as matrix elasticity, fluid shear stress, and tissue geometry could influence cell fate.^20–22^ Therefore we hypothesized that fluid flow modulates cell behavior and signaling in pancreatic progenitor cells.

In the present study, we used human iPSC-derived pancreatic progenitors to study the developing pancreas. The pancreatic progenitors were expanded following previously established protocols.^23^ These cells expressing PDX1, SOX9, and NKX6.1 are multipotent and are capable of differentiating into ductal, acinar, and endocrine cell types.^23,24^ To investigate the effect of physiological fluid flow, we developed a microfluidic platform compatible with conventional static culture and generation of mPa-level shear stresses. We analyzed gene expression changes and cell patch expansion patterns after fluid flow applied for 24 hours. Our findings establish pancreatic progenitor cells as shear stress-sensitive and help lay the foundation for understanding biophysical regulation of the developing pancreas.

## 2. Methods

### 2.1. Cell expansion

Pancreatic progenitors were differentiated from the hiPSC line CRTDi004-A (RRID:CVCL_YR23) (https://hpscreg.eu/cell-line/CRTDi004-A) and expanded (Fig. 1A) following a previously published protocol.^23^ Polystyrene cell culture 24-well plates (Sarstedt) were coated with 50 μg/mL of human fibronectin solution from plasma (Corning) for 30 min at 37°C to reach a theoretical surface concentration of 5 μg/cm^2^. Cryopreserved differentiated pancreatic epithelium (PE) cells^23^ at passage 7 (P7) from the CRTDi004-A (RRID:CVCL_YR23) line were thawed and expanded in 24-well plates. During expansion, the cells were plated at a density of 8.24 x 10^4^/cm^2^ in ePE (expanding PE) medium consisting of DMEM/F12 GlutaMAX (Fisher Scientific), 64 ng/mL FGF2 (Thermo Fisher Scientific), 1× B-27 (Fisher Scientific), 10 μM SB431542 (Selleck Chemicals), and 1% penicillin-streptomycin (Fisher Scientific). The medium was supplemented with 10 μM ROCK inhibitor Y-27632 (Selleck Chemicals) for the first 24 hours of each passage to reduce apoptosis and changed to ePE medium without ROCK inhibitor afterwards. The cells were then fed every 3-4 days. During serial expansion, cell stocks were cryopreserved at P9, P11, and P13. For shear stress experiments, cryopreserved cells at P11 or P13 were thawed, expanded for two or more passages, and seeded into the microfluidic devices between P15 and P17.

**Figure 1.**
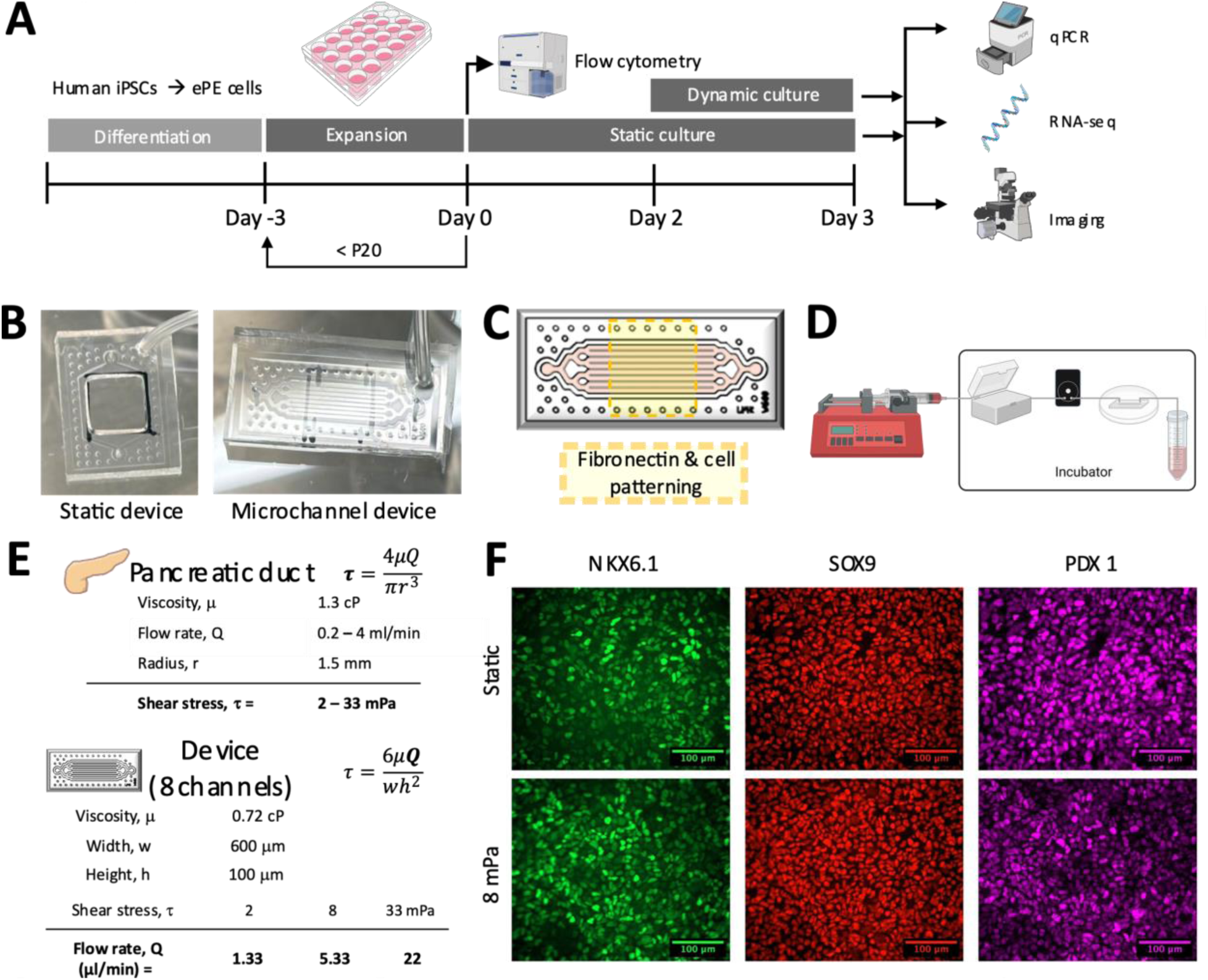
Laminar shear stress was applied to pancreatic progenitor cells using a custom designed microfluidic device. (A) Timeline of the experimental workflow. (B) Photos of static device and microchannel device vacuum sealed onto the dish substrate. (C) Schematic of the microchannel device showing selective patterning of protein and cells. (D) Schematic of the fluidic system containing syringe pumps, water bath, bubble traps, devices, and waste collection tubes, shown from left to right. (E) *In vitro* flow parameters were calculated to recapitulate physiological shear stresses in the adult pancreas, in lieu of available analogous embryonic values. (F) Representative immunostaining images of pancreatic progenitor cells show persistence of cell state marker expression after 24 hours of 8 mPa shear.

### 2.2. Shear stress calculation and device design

The microfluidic devices used in the present study incorporated vacuum-assisted reversible sealing.^25,26^ This feature enabled static culture in open well format before switching to shear stress application in microchannels (Fig. 1B). Wells or microchannels were designed with vacuum chambers in the surrounding area. Reversible sealing also made it possible to customize protein and cell surface patterning (Fig.1C). To facilitate easy device transfer between biosafety cabinet and incubator, syringes with stoppers^27^ were used as the vacuum source. Each fluidic system (Fig. 1D) was comprised of medium reservoirs (syringes mounted on a syringe pump), a water bath, bubble removers, microfluidic devices, and waste collection tubes.

Using the formula

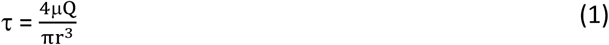

and previously reported values for pancreatic duct diameter,^28–31^ fluid viscosity,^32^ and flow rate,^33,34^ we estimated the shear stress in pancreatic duct to range from 2 to 33 mPa (Fig. 1E). Relatively few studies have explored cellular responses to mPa-level shear stresses.^11,35–37^ In our study, we chose 3 shear stress magnitudes representing low, medium and high magnitudes (LSS: 2 mPa; MSS: 8 mPa; HSS: 33 mPa) to cover the estimated range of physiological shear stress.

In order to generate all 3 shear stress magnitudes within the same perfusion device for 24 hours using readily available syringes (≤ 60mL) and standard syringe pumps, we designed channel dimensions to be 600 microns in width (w) and 100 microns in height (h). We calculated required flow rates (Q) using a medium viscosity (μ) of 0.72 cP and the formula for channels with rectangular cross sections:

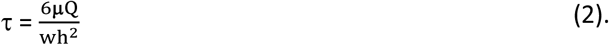

For each perfusion device containing 8 parallel channels, 24-hour perfusion at the 3 shear stress magnitudes required a minimum of 2mL, 8mL, and 32 mL of medium.

Human pluripotent-stem-cell-derived pancreatic progenitors have a relatively low oxygen uptake rate of approximately 0.016 pmol/s per 1500 cells.^38^ Based on an estimated 60,000 cells per microchannel, the total oxygen consumption was approximately 0.64 pmol/s. Applying a previously described oxygen mass balance considering convective oxygen delivery alone,^39^ a minimum flow rate of 0.5 μL/min per channel would be needed to balance the cellular oxygen consumption in devices made of gas-impermeable materials. The MSS and HSS flow rates of 0.67 and 2.75 μL/min exceeded this threshold, whereas the LSS flow rate of 0.17 μL/min did not. In practice, PDMS provides an additional route for oxygen delivery. The combined contributions of oxygen transport through PDMS and convective delivery via perfusion were expected to maintain adequate oxygen availability under all three shear stress conditions (Supplemental method).

### 2.3. Microfabrication

Film photomasks were designed in AutoCAD and printed (JD Photo Data, UK). Device molds were made following photolithography procedures including SU-8 negative photoresist spin coating on silicon wafer, pre-baking, UV exposure, post-baking, development, hard baking, and structure depth measurement with Profilometer. Device molds were then used to make microfluidic devices cast in poly(dimethylsiloxane) (PDMS, Sylgard 184, Darwin Microfluidics).

### 2.4. Static and dynamic culture in microfluidic devices

Static PDMS well devices were vacuum sealed onto BioLite 60mm dishes (Thermo Fisher Scientific) to enable selective coating of the central regions (1×1cm^2^) with 5 μg/cm^2^ of fibronectin. Following 30 min incubation at 37 °C, fibronectin solution was removed. Next, ePE cells detached from each 24-well were seeded into 4 PDMS well devices in ePE medium supplemented with ROCK inhibitor, which was changed to ePE medium without ROCK inhibitor 24 hours later. After 48 hours of static culture to reach confluence, ePE medium exchange was performed for static control conditions. For paired shear stress conditions, cells were briefly submerged under basal medium (DMEM/F12 GlutaMAX) while the PDMS well devices were replaced with parallel-channel devices, to avoid trapping bubbles in the channels, inlets, and outlets. Syringe pumps (PHD ULTRA, Harvard Apparatus; AL-4000, World Precision Instruments) were used to set flow rates to achieve desired shear stress. After priming the fluidic system with ePE medium, tubing was connected to the inlet and outlet of parallel-channel devices. Experiments were performed using paired static and shear stress samples.

### 2.5. Flow cytometry

Cells were dissociated from 24 well plates, fixed in 4% formaldehyde for 15 min, and washed with 1x PBS. Fixed cells were permeabilized with 0.3% Triton X-100 in PBS containing 5% donkey serum and 0.5% w/v IgG-free BSA for 30 minutes at room temperature. Permeabilized cells were then incubated with primary antibodies against PDX1 (1:800), SOX9 (1:1000), and NKX6.1 (1:100) in 0.5% w/v IgG-free Bovine Serum Albumin (BSA) in PBS overnight at 4 °C. After PBS washes, cells were incubated with secondary antibodies conjugated with fluorophores (1:1000) for 1 hour at room temperature. Antibody and reagent sources are indicated in Table S1. Cells were analyzed using an LSR Fortessa (BD Biosciences), and data were analyzed with the FACS Diva version 9.0.1 (BD Biosciences).

### 2.6. Immunofluorescence staining and microscopy

For immunostaining experiments, PDMS devices were released from vacuum and removed from the dishes after 24 hours static or dynamic culture. Cells were washed with 1x PBS and fixed with 4% formaldehyde, followed by washes in PBS. After permeabilization in PBS with 0.1% Triton X-100 for 15 minutes, samples were treated with blocking solution consisting of 0.3% Triton X-100 and 5% normal donkey serum in PBS. Samples were incubated with primary antibodies against PDX1 (1:800), SOX9 (1:1000), and NKX6.1 (1:100) in blocking solution for two hours at room temperature. Cells were then washed with PBS and incubated with secondary antibodies conjugated with fluorophores (1:1000) and Hoechst (1:3500) in blocking solution for one hour at room temperature. Samples were imaged using a widefield microscope (Axiovert 5, Carl Zeiss) with 40x objective (LD A-Plan NA=0.55).

For experiments staining YAP1 and phosphorylated YAP1 (p-YAP1), cells were fixed with 4% formaldehyde on the chip prior to device detachment and PBS washes. Following permeabilization in PBS with 0.2% Triton X-100 for 5 minutes, samples were treated with blocking solution consisting of 2% IgG-free BSA, 0.3% Triton X-100 and 5% Goat Serum in PBS. Samples were incubated with primary antibodies against YAP1 (1:200) and p-YAP1 (1:100) in blocking solution overnight at 4°C. Cells were then washed with PBS, incubated with secondary antibodies conjugated with fluorophores (1:1000) in blocking solution for 2 hours at room temperature, followed by incubation with Hoechst (1:3500) in blocking solution for 10 minutes at room temperature. Samples were imaged on a confocal laser scanning microscope (Axio Observer.Z1/7 with LSM980, Carl Zeiss) using a 63x objective (C Plan-Apochromat, NA=1.40) and oil immersion.

### 2.7. Generation of putative shear-responsive genes expressed in fetal pancreas

Titles and abstracts of shear stress-related PubMed-indexed articles were screened for mentions of genes and proteins using a previously published protocol.^40^ Applying an article count ≥10 filter generated 212 genes frequently mentioned in shear stress literature agnostic to organ system context. A previously published single cell transcriptomic dataset^41^ was used to verify the expression of the shear stress-associated genes in fetal pancreas. Based on the expression patterns, 13 putative shear-responsive genes and a progenitor marker gene *NKX6.1* were selected for qPCR measurement.

### 2.8. RNA processing

At the 24-hour time point following the initiation of the shear stress application, cells were lysed directly on the chip using Lysis Buffer (Invitrogen) supplemented with 2-Mercaptoethanol. Total RNA was extracted using Invitrogen™ PureLink™ RNA Micro Scale Kit (Fisher Scientific) following manufacturer’s instructions. Samples with low A_260_/A_230_ absorbance ratios were purified via precipitation with sodium acetate (Fisher Scientific) and reconstitution in nuclease-free water prior to RNA sequencing.

### 2.9. RT-qPCR

Total RNA was reverse transcribed using an Applied Biosystems^TM^ High-Capacity cDNA Reverse Transcription Kit (Thermo Fisher Scientific). qPCR was carried out with PowerTrack™ SYBR Green Master Mix (Thermo Fisher Scientific), and all samples were tested in duplicates. mRNA levels of queried genes were normalized to the average of static control samples. Primers for all genes are listed in Table S2.

### 2.10. RNA-sequencing and analysis

RNA samples for sequencing had concentrations >10 ng/μl and RIN scores >9 (Agilent Bioanalyzer), indicating sufficient concentration and quality of RNA, respectively. cDNA libraries were prepared using poly-dT magnetic bead enrichment of messenger RNA, fragmentation, and first strand synthesis using random hexamer primers, followed by the second strand cDNA synthesis using either dTTP for non-strand specific library (Novogene). Samples were analyzed as 150 bp paired end reads using a NovaSeq X Plus Series (Illumina) sequencer. Reads were processed by Novogene for QC (GC content, sequence error rate, and removal of low quality reads) before alignment using HISAT2^42^ to the human reference genomic chg38. A total of 35 samples, comprised of 31 PDMS-device samples culture under static or shear stress conditions and 4 well-plate samples, were sequenced. Sequencing produced 13.8 ± 1.0 GB (avg. ± st. dev.) sequencing data per sample with 86.2 ± 2.9% of paired reads simultaneously aligned to the genome (‘proper reads’) per sample. Downstream RNA-seq analysis focused on the 31 PDMS-device samples. Gene read counts per Ensemble ID and per sample (Table S3) were downloaded from NovoMagic Gene Expression Analysis Quantification. Read counts were pre-processed to remove low count genes (<10 counts/sample in 27 or more samples). Remaining genes (those with counts ≥10 in at least 5 samples) were referred to as expressed genes in this study. DESeq2^43^ was used to analyze bulk RNA-seq data with batch effects modeled in. Pairwise contrasts were performed between the shear stress samples and all 14 static samples. Effect size shrinkage was performed using Apeglm.^44^ Differentially expressed genes (DEGs) were selected based on |Log_2_FC shrunk|>1 and P_adj_<0.05. For principal component analysis and gene expression visualization using variance stabilized transformed data (vsd), batch effects were removed using Limma.^45^ One-way ANOVA with Games-Howell post hoc test was performed for the union list of differentially expressed genes to identify genes that varied between static and any shear condition.

### 2.11. Functional enrichment analysis

Gene Set Enrichment Analysis (GSEA) was performed using Webgestalt^46^ on the expressed genes ranked by Wald stat. Default parameter settings were used with the exception of significance level: instead of top 10 results, all results with FDR<0.05 were shown. The top 5 Hallmark gene sets were then selected by applying a |Normalized Enrichment Score|≥2 filter. The ranked gene lists and the other statistically significant Hallmark gene sets can be found in Table S4. The leading edge gene sets can be found in Supplementary Data.

Gene Ontology (GO) overrepresentation analysis (ORA) was performed using g:Profiler^47^ on differentially expressed genes. Default parameter settings were used with the exception of the statistical domain scope, where all expressed genes (after prefiltering) were used as the background gene list with the option “custom over annotated genes” chosen. To identify top GO terms, two filters were imposed: (1) term size ≤ 2000 and (2) P_adj_ ≤ 0.005. To calculate DEG similarity among GO terms within each of the 3 GO categories, Jaccard and Overlap Combined Coefficient (50%/50%) was used to generate binder plots. The complete GO ORA results can be found in Table S5.

### 2.12. Image analysis

Fiji was used for analyzing images of YAP1 and p-YAP1 staining following recommended procedures.^48^ Briefly, background subtraction (sliding paraboloid, radius 100 pixels) was applied to images before analysis. Nuclear staining images were converted to binary masks following threshold adjustment. Mean fluorescence intensity (FI) of YAP1 and p-YAP1 inside and outside the nuclear masks was measured to enable comparison between static and shear stress samples. Nuclear-to-cytoplasmic YAP1 mean FI ratios were calculated to quantify the extent of YAP1 nuclear exclusion.

### 2.13. Barrier-based migration assay under shear stress

Fibronectin was initially coated in a 1×1 cm^2^ region using a square-well PDMS device vacuum-sealed onto a 60mm cell culture dish. This was then replaced with a narrow-well PDMS device to culture cells in the central region (0.7×1 cm^2^). Once cells reached confluence after 2 days of static culture, the well device was removed to permit lateral expansion of the cells into fibronectin-coated cell free zones. For shear stress conditions, the well device was replaced with a parallel-channel device sealed using a vacuum pump. Devices were then transferred to an Ibidi stage-top incubator attached to a widefield microscope (Axio Observer.Z1/7, Carl Zeiss). The system was equipped with temperature control and a CO_2_ humidity chamber at 37°C and 5% CO_2_ throughout the experiment. A heat block was maintained at 37°C to warm up medium in the tubing right before its entry into the parallel-channel device. The experiment was performed using a 20x objective (LD Plan-Neofluar, NA=0.4). Phase images of cell patch boundaries were taken every 6 hours over the course of 24 hours. The change in cell-covered area (*ΔA*) was measured (Table S6), and the average net displacement distance was calculated based on *D=ΔA/w* with *w* being the cell patch width within the field of view.

### 2.14. Statistical analysis

Data visualization and statistical analyses were performed using GraphPad Prism, R and Python. N refers to number of replicates, whereas n refers to number of experiments. Statistical significance (p<0.05) was calculated using Tukey’s multiple comparison, Games-Howell multiple comparison.

## Results and Discussion

### 3.1. Custom-designed devices for applying 24-hour mPa-level shear stress at three magnitudes

Chips intended for cell culture typically have large channel heights to ensure sufficient internal volume for nutrient supply and waste dilution. However, generating the same shear stress in a taller channel requires a higher flow rate and increased media consumption; this can be particularly costly for stem cell and progenitor cell culture. To mitigate this, recirculation systems are often utilized for long-term perfusion. In this study, we aimed to apply physiologically relevant levels of shear stress (LSS: 2 mPa; MSS: 8 mPa; HSS: 33 mPa) to pancreatic progenitor cells for 24 hours using a single perfusion device design, with infusion achieved via cost-effective options such as syringe pumps. Therefore, we designed devices with vacuum chamber features compatible with reversible sealing. This enabled static culture in open-well format followed by shear stress application in microchannels. A previously established protocol for pancreatic progenitor cell culture in well plates could thus be directly applied to PDMS wells, eliminating the need to adapt protein coating, cell seeding, and media exchange procedures required for static culture in microchannels. In these conditions, the cells were oriented with their basal side facing the fibronectin coating and their apical side facing the fluid, as *in vivo*. Reversible sealing also permitted a lower microchannel height (100 μm), which kept daily media consumption per device below 10 mL for LSS, 20 mL for MSS, and 40 mL for HSS, including the additional volume required for fluidic system priming. For 24-hour perfusion experiments, standard syringes (≤60 ml) and syringe pumps were therefore sufficient. To better model lumenal lining in developing pancreas, cells were allowed to reach confluence prior to shear stress application.

Perfusion devices were designed with constant channel width (600 μm) and height (100 μm), from inlet to outlet. As fluid diverges into parallel channels, the flow rate per channel and wall shear stress are both reduced by half at each bifurcation. To ensure exposure to uniform shear stress, cells were patterned within the 8-channel region. Pancreatic progenitor cells expanded in well plates expressed progenitor cell markers PDX1, SOX9, and NKX6.1 (Fig. S1). After 24 hours of shear stress exposure, the markers remained detectable (Fig. 1F).

### 3.2. Pancreatic progenitor cells respond to mPa-level shear stresses

Next, we investigated whether pancreatic progenitor cells undergo gene expression changes in response to the physiologically relevant mPa-level shear stresses. Experiments were conducted using paired static and shear stress samples. Utilizing Gene Frequency Analzyer^40^ and existing transcriptomic datasets^41^ (Fig. 2A), we generated a short list of putative shear stress-responsive genes that were frequently cited in the shear stress literature and also expressed in human fetal pancreas. qPCR results (Fig. 2B, Fig. S2) demonstrated that genes such as *FOS* and *EGR1* (Fig. 2Bi-iv) had statistically significant expression changes at HSS (33 mPa), whereas *CD44* and *TFPI* (Fig. 2Bv,vi) had statistically significant expression changes at all 3 shear stress levels (2, 8, 33 mPa). Within the range of shear stresses examined, qPCR results suggested that gene expression responses to applied shear stress could be either magnitude-dependent or magnitude-independent. These patterns could result from distinct mechanosensing and mechanosignaling pathways, with different activation thresholds and dynamic ranges. *ACTB*, which is commonly used as a housekeeping gene, was found to be upregulated by HSS (Fig. 2bvii), similar to its behavior in renal epithelial cells.^11^ We then quantified expression of progenitor cell state maker *NKX6.1*, as it is known to be influenced by cell culture conditions and passage numbers.^23^ However, we detected no significant changes in *NKX6.1* levels (Fig.S2J). *KRT19*, one of the ductal marker genes, was found to be upregulated at HSS (Fig. 2Bviii).

**Figure 2.**
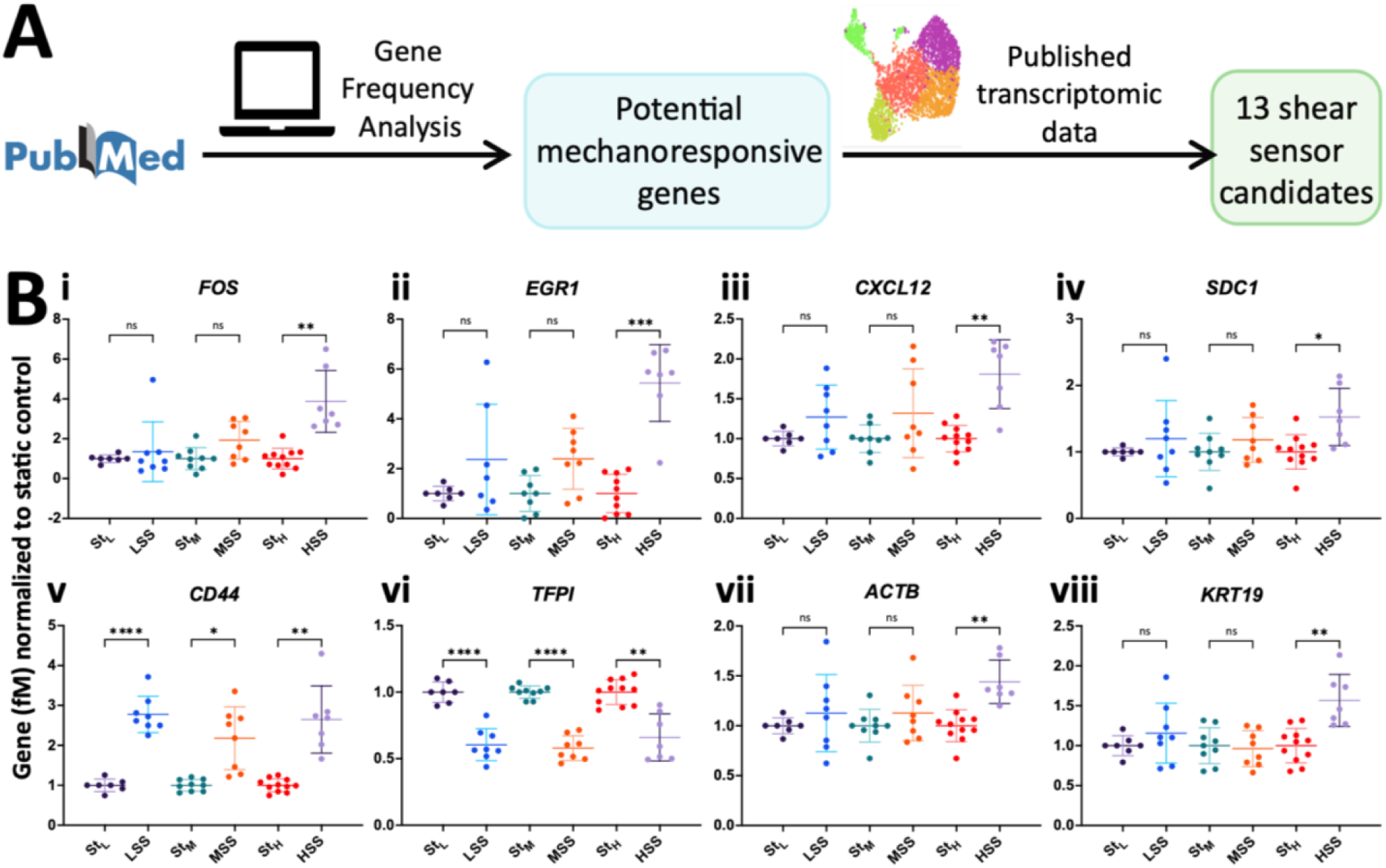
Pancreatic progenitor cells responded to 24 hours of mPa-level shear stress with gene expression changes. (A) schematic of workflow for generating a short list of data-mined^40^ putative shear stress-responsive candidates for qPCR measurement. (B) Scattered dot plots of shear stress-responsive genes from qPCR experiments. Non-responsive genes can be found in Figure S2. N≥7 replicates per condition, n=4 independent experiments per SS-vs-Static comparison. Brown-Forsythe and Welch ANOVA with Games-Howell post hoc test. Abbreviations: St = Static, L/LSS = low shear stress, M/MSS = medium shear stress, H/HSS = high shear stress. ns: P>0.05, *: P≤0.05, **: P≤0.01, ***: P≤0.001, ****: P≤0.0001. The 7 unresponsive genes are shown in Figure S2D-J.

### 3.3 Shear stress-induced transcriptome-wide changes

In order to understand the transcriptome-wide shear stress effect on pancreatic progenitor cells, we performed bulk RNA-seq. Experiments were conducted to include paired static and shear stress samples (5 LSS, 6 MSS, 6 HSS, and 14 static), with a few experiments including more than 1 shear stress level. In total, among the 31 samples from shear stress experiments, 20,950 genes satisfied the expression criterion (≥10 counts in at ≥ 5 samples). Batch effects were present (Fig.S3A). After batch effect removal (Fig.S3B-F), Principal Component Analysis results (Fig. 3A) showed that shear stress samples were well separated from static samples along PC1. While the shear stress samples were largely separated along PC2, some degree of intermixing was observed, especially between MSS and HSS samples.

**Figure 3.**
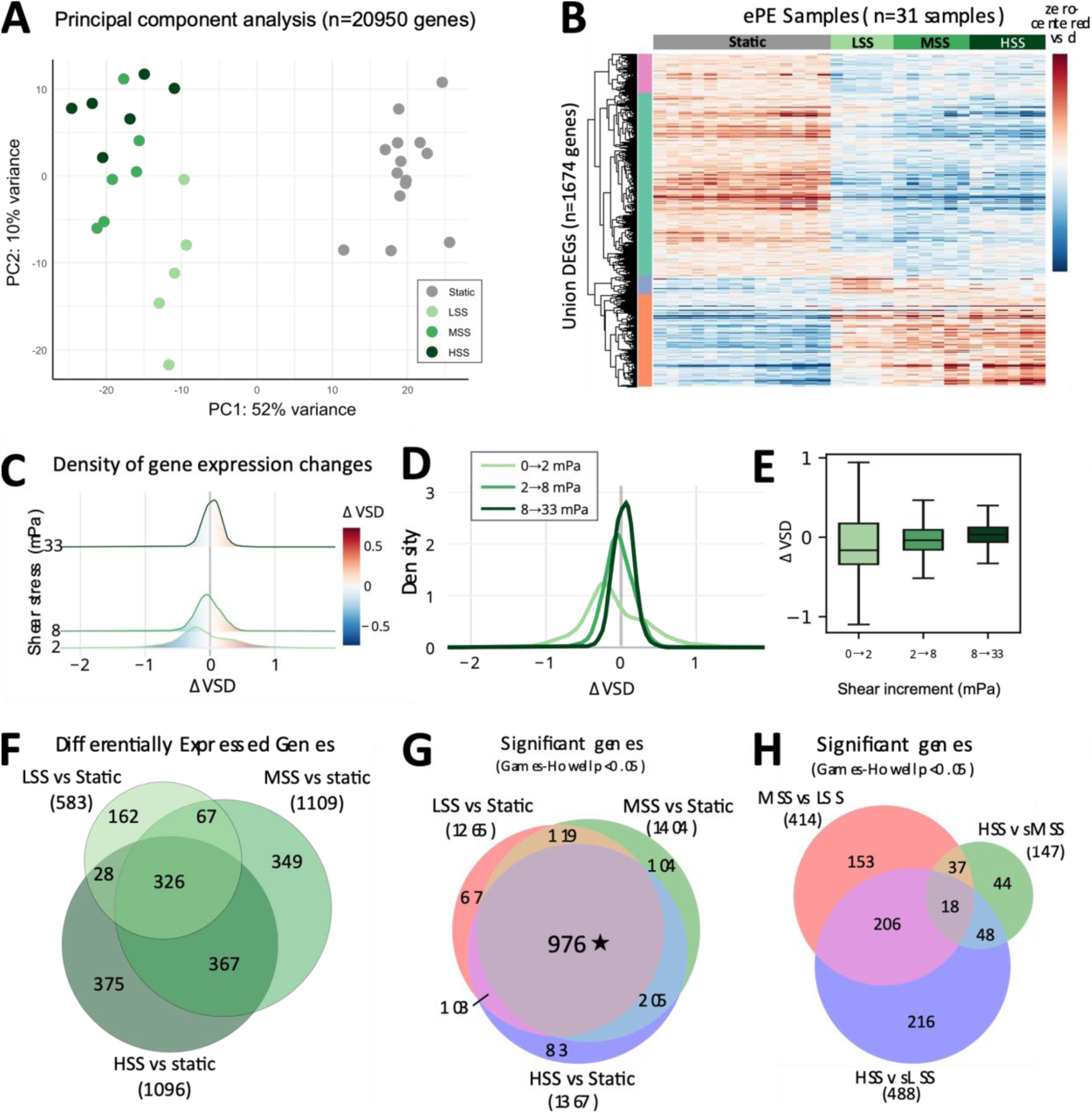
24 hours of laminar shear stress at (2, 8, 33 mPa) induced transcriptome-wide changes in pancreatic progenitor cells. (A) Principal component analysis of 31 samples based on the 20,950 expressed genes after batch effect correction. (B) Hierarchically clustered heatmap of differentially expressed genes across conditions and samples. (C) Ridgeline plot, (D) overlay line graph, and (E) Tukey boxplots showing distributions of gene expression changes at each shear stress increment. (F) Venn diagram showing overlap of shear-responsive DEGs. (G, H) Venn diagrams comparing the gene subsets that showed statistical significance between conditions based on one-way ANOVA performed on 1674 DEGs followed by Games-Howell post hoc tests. N≥5 replicates per condition, n≥3 experiments per SS-vs-Static comparison.

Pairwise contrasts using DESeq2 showed results broadly consistent with the RT-qPCR measurements for the data-mined shear-responsive genes (Fig. S3G). Differential expression analysis identified a union of 1674 differentially expressed genes (DEGs) out of the 20,950 genes expressed in ePE cells (Table S3).

Among these 1,674 union DEGs, more genes were downregulated (N=1,012) than upregulated (N=662) upon shear stress exposure (Fig.3B). While subsets of genes (pink and blue clusters) showed biphasic responses, the majority of the genes (turquoise and orange clusters) exhibited predominantly monotonic responses with increasing shear stress magnitude.

The largest transcriptomic response occurred between 0 and 2 mPa, despite this being the smallest differential in applied shear stress. The 0-to-2 mPa transition (Fig. 3C-E) had a broader, flatter Δvsd distribution spread and the largest interquartile range (IQR=0.51), suggesting that a greater proportion of genes underwent substantial expression shifts. The median value (-0.16) of the 0-to-2 mPa distribution and its dominant peak in the negative range again demonstrated that more genes were downregulated. At higher shear stresses (8 and 33mPa), the peak and the median of the Δvsd distribution shifted rightwards relative to 0-to-2 mPa. The 2-to-8 mPa increment elicited greater expression changes than the larger shear stress increment of 8-to-33 mPa (wider Δvsd spread, larger IQR: 0.25 vs 0.18). Overall, gene expression changes started to plateau as shear stress increases, within the range tested.

Of 1,674 differentially expressed genes (Fig.3F), more genes met the DEG criteria (|Log_2_FC shrunk|>=1 and P_adj_ <0.05) for MSS-vs-static and HSS-vs-static comparisons than LSS-vs-static. To understand whether this was due to difference in effect size, we performed Welch ANOVA followed by Games-Howell post hoc tests (Table S3) on all DEGs. The apparent difference in the number of DEGs was largely driven by the effect-size threshold. This analysis revealed that the number of genes with statistically significant gene expression changes were more similar between static and shear stress conditions than the number of DEGs implied. LSS at 2 mPa appeared sufficient to invoke a response from 1,265 genes out of the 1,674 union DEGs tested (Fig. 3G), despite only 583 DEGs in LSS-vs-static. Compared to static conditions, 976 DEGs responded to all 3 shear stress magnitudes.

When inter-shear stress comparisons were performed (Fig. 3H), over 400 genes had statistically significant differences for both MSS-vs-LSS and HSS-vs-LSS, while only 147 genes were significantly different between MSS and HSS. 18 genes showed statistically significant differences among all 3 shear stress levels. 206 genes differed significantly for both MSS-vs-LSS and HSS-vs-LSS, but not between MSS and HSS, suggesting their gene expression may plateau above 8 mPa.

### 3.4 Transcriptomic responses to shear stress are associated with cell cycle, endocrine differentiation, and receptor-ligand signaling

To understand the functions associated with global transcriptional changes, we first performed Gene Set Enrichment Analysis (GSEA) using the 20,950 expressed genes. This analysis revealed E2F targets, MYC targets, G2/M checkpoint, mTORC1 signaling, and pancreatic β-cell as the top 5 most enriched Hallmark gene sets^49^ (Fig. 4A, Fig. S4A). These terms had the highest average |NES| values, with absolute value Normalized Enrichment Score (NES) ≥2 for all 3 pairwise comparisons.

**Figure 4.**
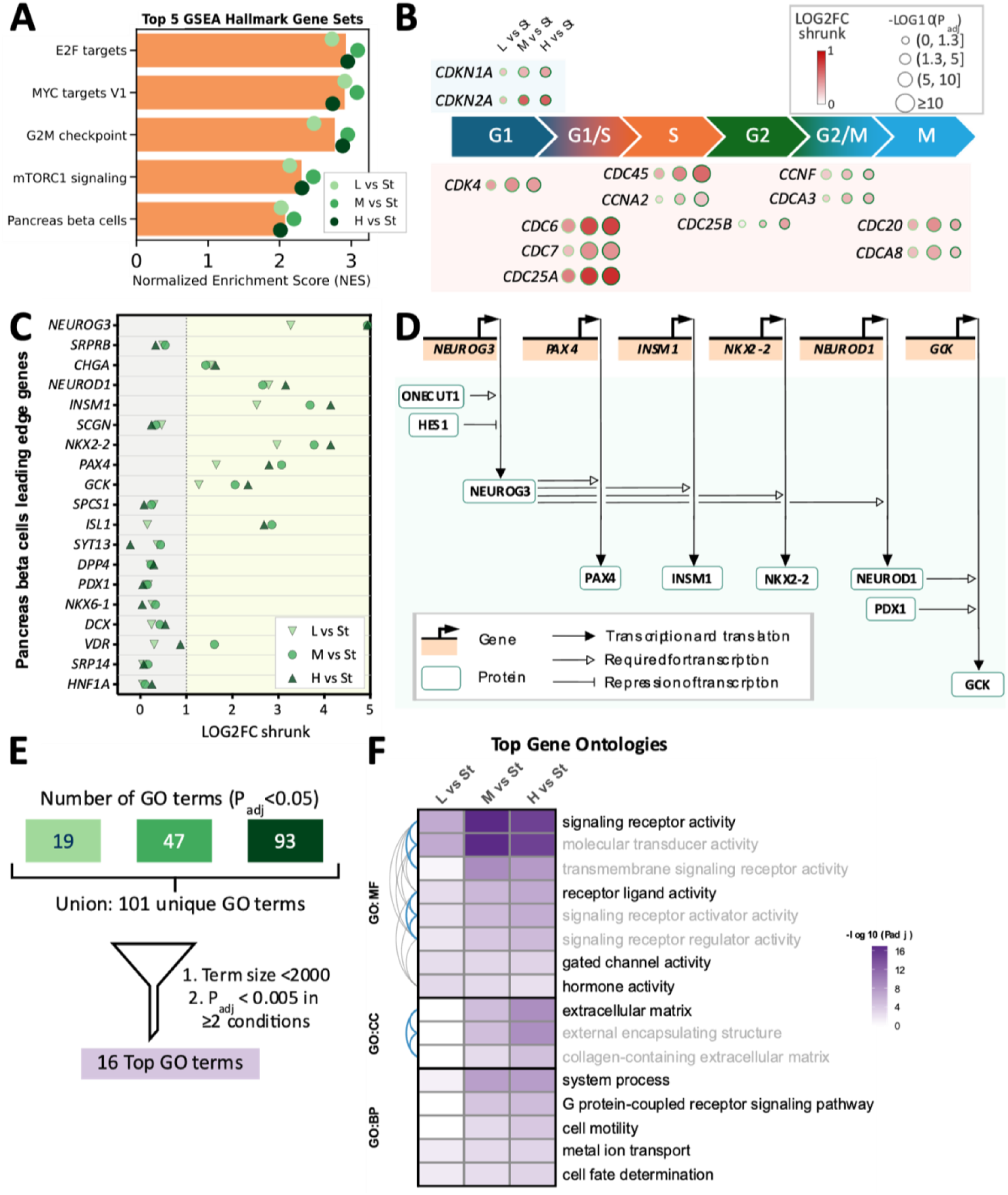
Pathways associated with shear-induced gene expression changes. (A) Bar graph showing average Normalized Enrichment Scores of the top 5 Hallmark gene sets from GSEA. (B) Dot heatmaps of leading-edge genes associated with cell division cycle and cyclins. (C) Dot plot of leading-edge genes from the pancreatic beta-cell Hallmark gene set. (D) Reduced Reactome pathway diagram (Regulation of beta cell development)^60^ highlighting NEUROG3 as a trigger of the endocrine differentiation transcriptional program. (E) Flow chart showing the selection process of top GO terms. (F) Heatmap of the top overrepresented GO terms, with binder plot connectors indicating gene set similarity (gray: 0.6 ≤ combined Jaccard-overlap coefficient <0.7; blue: ≥ 0.7) within molecular function, cellular component, and biological process categories. Within each group of similar GO terms, the first-listed term is shown in black and the remaining terms in gray.

The top 4 Hallmark leading edge gene sets included cyclin- and cell division control-related genes that are primarily associated with different phases of cell cycle (Fig. 4B). Surveying of KEGG pathway also helped identify additional cell cycle genes affected by fluid shear stress (Table S7), among which 4 genes (*CCNA1*, *CDKN2B*, *SFN*, *GADD45G*) met DEG criteria in at least 1 shear stress vs static pairwise comparison (Fig. S4B). Previously the 3 cell cycle arrest molecules (*CDKN2B*, *SFN*, and *GADD45*) were also reported to be upregulated in renal epithelial cells exposed to mPa fluid shear stress.^11^

In contrast to the cell cycle gene sets, the enrichment of pancreas beta cell Hallmark gene set is cell type-specific. The leading-edge genes included *CHGA*, encoding the pan-endocrine marker chromogranin A, and *NEUROG3*, encoding the master regulator^50^ of endocrine differentiation neurogenin-3 (Fig. 4C). All 3 shear stress levels resulted in around 3-fold increases in *CHGA* expression. Compared to static conditions, exposure to LSS led to an approximately 10-fold increase in *NEUROG3* expression, whereas MSS and HSS resulted in 30-fold increases. NEUROG3 is widely considered to be the master regulator that initiates endocrine differentiation^51–53;^ its activity is required for transcription of other endocrine markers such as *NEUROD1*, *PAX4*, *INSM1*, *NKX2-2*, and *GCK* (Fig. 4D). The upregulation in these genes (Fig. 4C, Figure S4C) is consistent with induction of *NEUROG3* transcription and indicative of increased NEUROG3 protein activity under flow. This result suggests fluid shear stress may influence genes involved in cell fate decision making. In adult pancreas, mature endocrine cells are not in direct contact with the lumenal secretion. However, *NEUROG3*-expressing cells have been reported to delaminate from the ductal structures during pancreatic development.^54^ Therefore, fluid flow has the potential to modulate pancreatic endocrine differentiation process during embryonic development.

In addition to upregulating the top 5 Hallmark gene sets, fluid flow also downregulated gene sets related to inflammatory responses, interferon alpha and interferon gamma responses (Table S4). The anti-inflammatory^55^ and anti-proliferative^56^ effects of laminar shear stress are well-established in the field of vascular mechanobiology. While fluid shear stress appeared to also exert anti-inflammation effect on pancreatic progenitor cells, further research is needed to elucidate the net effect of shear stress on proliferation, as both cell cycle progression-related gene sets and cell cycle arrest genes showed increased expression. Fluid flow also upregulated oxidative phosphorylation gene set and downregulated hypoxia gene set for pancreatic progenitor cells (Table S4). Previous studies have reported that that laminar shear stress may shift endothelial cells from glycolysis to oxidative phosphorylation,^57^ inhibit HIF-1a signaling,^58^ and downregulate hypoxia-related genes^59^ compared to static culture. Our data suggests that similar shear-induced responses might occur in pancreatic progenitor cells as well.

To identify processes associated with the genes highly responsive to shear stress, we performed Gene Ontology^29,61^ overrepresentation analyses on DEG gene lists using g:Profiler^47^ (Fig. 4E-F, Fig. S4D). This analysis additionally revealed expression changes in genes related to receptor-ligand activity, extracellular matrix remodeling, GPCR signaling, metal ion transport, and cell fate determination. Among the molecules connected to the overrepresented GOs, 14 gene families with ≥4 members were affected by fluid flow (Table S5). In particular, prostaglandin receptors were consistently upregulated in shear stress conditions. Gene expression of ion channels, including K^+^, Na^+^, Ca^2+^, and Transient Receptor Potential channels, were also modulated by shear stress. Overall, these findings suggest that shear stress elicits remodeling of diverse cell signaling processes.

### 3.5 Fluid shear stress triggers upregulation of *DLL1* and *DLL3*, accompanied by reduced YAP1 activity

Among the genes most strongly induced by shear stress were the Notch ligand genes *DLL1* and *DLL3* (Fig. 5A, Fig. S5A-B), both of which have been implicated in the regulation of Notch signaling. Relative to static controls, *DLL1* gene expression increased to 2.7-, 4.7-, and 6.4-fold, whereas *DLL3* gene expression increased to 7.3-, 15.8-, and 22.6-fold upon exposure to LSS, MSS, and HSS. DLL1 protein is known to act as trans-activator of NOTCH1 receptor and a key signaling molecule during endocrine differentiation. DLL1—NOTCH1 protein interaction leads to lateral inhibition, resulting in salt and pepper patterns of NEUROG3^+^ endocrine fated cells and NEUROG3^-^ cells during development.^62^ This raises the possibility of cell subpopulations after shear stress application. Unlike DLL1, DLL3 protein functions as a cis-inhibitor sequestering NOTCH1 in the Golgi leading to Notch inhibition. *DLL1* and to a lesser extent *DLL3* are direct targets of the Notch-activated transcription factor HES1 and are activated upon HES1 down-regulation.^63^ In parallel to NEUROG3 activation, *DLL1* is also a direct target of NEUROG3 and NEUROG3 is not necessary for its expression.^52,64^ While *DLL3* gene expression can be detected in endocrine precursor cells and beta cells according to the previously published fetal pancreas transcriptomic dataset,^41^ very few studies have shed light on the role of *DLL3* in pancreatic development. In contrast, DLL3 has recently attracted considerable attention in cancer research concerning neuroendocrine neoplasms, ^65–67^ and it has been reported to modulate DLL1—Notch signaling.^68^

**Figure 5.**
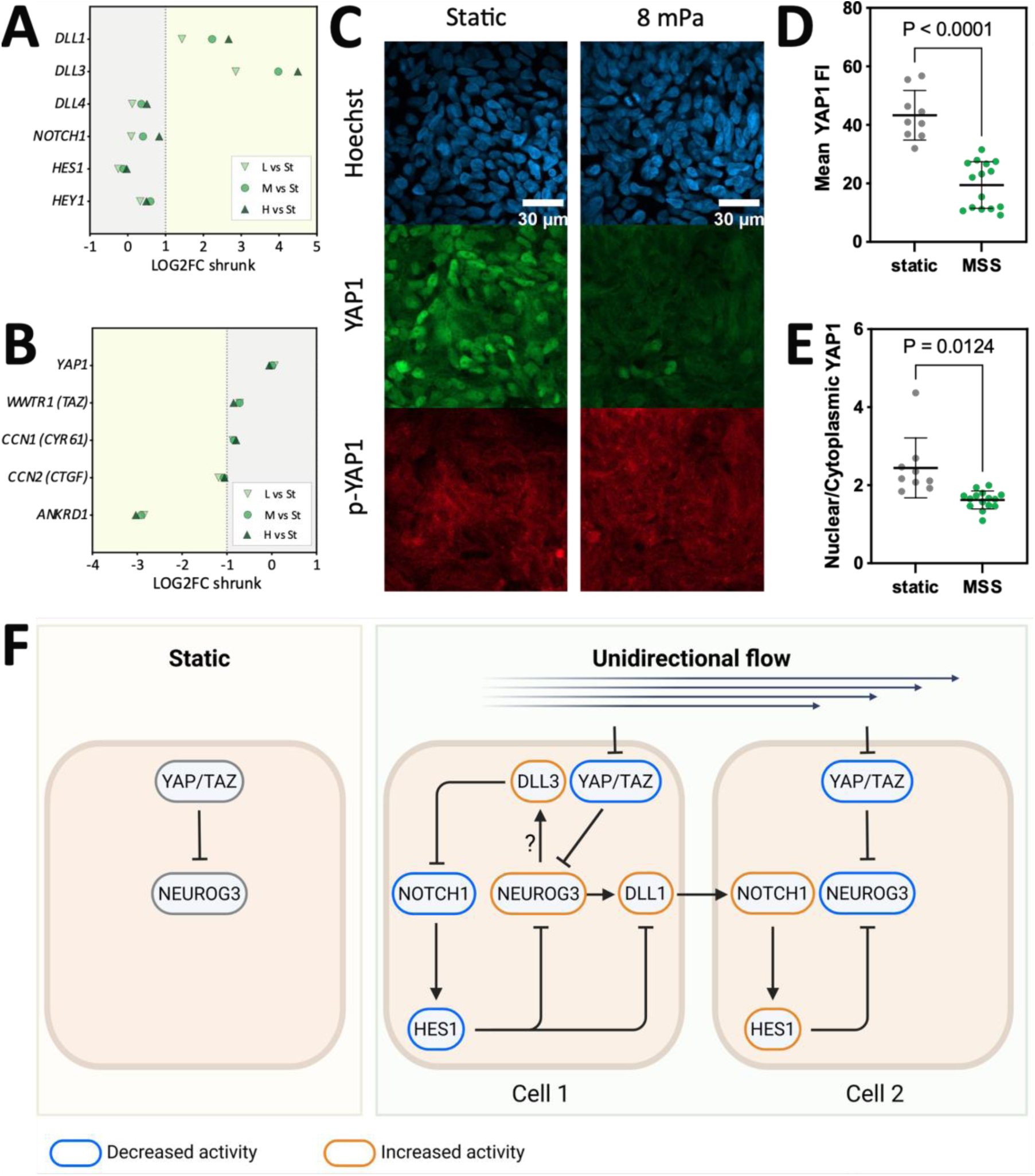
Mechanosensitive signaling pathways in response to fluid shear stress. (A) Dot plot shows expression changes of Notch signaling genes and HES1 target genes after 24 hours of fluid flow exposure. (B) Dot plot shows gene expression changes for *YAP1*, *WWTR1*, and YAP/TAZ target genes. (C) Representative confocal images of pancreatic progenitor cells stained for nuclei, YAP1, and phosphorylated YAP1 (Ser127) after 24 hours of MSS. (D-E) Quantification of mean YAP1 Fluorescence Intensity (FI) and ratio between nuclear and cytoplasmic mean YAP1 FI. N≥9 replicates, n=3 experiments. Welch’s t-test. (F) Schematic of the proposed model showing YAP1-mediated *NEUROG3* modulation in response to unidirectional fluid flow.

We also observed a statistically significant increase in gene expression (to 1.4-fold under HSS) for *DLL4*, which encodes another NOTCH1 trans-activating ligand (Fig. 5A, Fig. S5C). *NOTCH1* gene expression also increased modestly (Fig. 5A, Fig. S5D), although the magnitude of this change was below the DEG cutoff of |Log_2_FC shrunk|=1. The Notch signaling target gene *HES1*^69^ did not show statistically significant expression changes based on our bulk RNA-seq results (Fig. 5A, Fig. S5E). However, spatially heterogeneous *HES1* expression resulting from DLL1—NOTCH1-mediated lateral inhibition may be obscured in bulk transcriptomic measurements. A previous study identified *DLL1*, *DLL3*, *DLL4*, and *HEY1* as HES1 target genes in pancreatic progenitors.^63^ Besides *DLL1, DLL3* and *DLL4*, *HEY1* expression also increased significantly to 1.5- and 1.4-fold under MSS and HSS, respectively (Fig. 5A, Fig. S5F). Beyond ligand-mediated regulation, high fluid shear stress (>1 Pa) has been reported to directly activate NOTCH1 signaling in endothelial cells^70^, but it is unclear whether mPa-level shear stress is sufficient to trigger NOTCH1 activation in pancreatic progenitor cells. However, given the established role of canonical Notch signaling in maintaining pancreatic progenitor identity and suppressing endocrine differentiation,^71^ NOTCH1 activation by shear stress, if present, would be expected to inhibit *NEUROG3* gene expression. Therefore, direct activation of NOTCH1 via HES1 by fluid flow is unlikely to be responsible for *NEUROG3* upregulation observed in the present study.

Recently, YAP1 has emerged as another key regulator of endocrine differentiation in pancreatic progenitor cells.^72–75^ YAP1 activity is also known to be influenced by various physical cues including fluid shear stress.

However, whether mPa-level shear stress can regulate YAP1 in pancreatic progenitor cells has not been investigated before. Previous studies have reported both activation and inhibition of YAP1 by fluid shear stress. These seemingly contradictory findings can largely be explained by study differences in species, cell type, fluid flow directionality, duration, and shear stress magnitude. In general, oscillatory flow has frequently been associated with increased YAP1 activity, whereas sustained unidirectional flow often promotes YAP1 cytoplasmic localization and reduced activity.^76,77^ Several studies have characterized biphasic temporal responses to unidirectional fluid flow, with YAP1 activity initially increasing shortly after flow onset before declining during prolonged exposure.^78,79^ In kidney epithelial cells, constant physiological shear stress at 100 mPa for 48 hours reduced YAP1 activity relative to the static culture, whereas 24 hours of 100 mPa followed by another 24 hours of constant pathological shear stress at 400 mPa reverted YAP1 activity to the static level.^80^ In our experiments, *YAP1* gene expression (Fig.5B, Fig. S5G) was not altered by 24 hours of constant MSS exposure (8 mPa), whereas *WWTR1* transcript levels (Fig.5B, Fig. S5H) were significantly reduced by approximately 40% under all 3 shear stress conditions. We also detected statistically significant downregulation of common YAP1/WWTR1 (YAP/TAZ) target genes such as *CCN1* (*CYR61*), *CCN2* (*CTGF*), and *ANKRD1* at all 3 shear stress levels (Fig. 5B, Fig. S5I-K). Immunofluorescence microscopy results (Fig.5C) showed a reduction in mean YAP1 fluorescence intensity (Fig.5D) and a decrease in nuclear-to-cytoplasmic YAP1 ratio under MSS (Fig. 5E). In contrast, phosphorylated YAP1 mean FI (Fig. S5L) showed no difference, while the p-YAP1:YAP1 ratio (Fig. S5M) was higher under MSS only due to the reduced mean YAP1 FI. Although phosphorylated YAP1 abundance was similar between static and MSS conditions at the end timepoint, reduced YAP1 abundance, decreased YAP1 nuclear localization, and downregulation of YAP/TAZ target genes together indicate diminished YAP1 activity after 24 hours of constant shear stress at 8mPa.

Given that 2-33 mPa shear stress upregulated endocrine-associated genes and reduced YAP1 activity, we propose a model in which unidirectional fluid flow promotes *NEUROG3* expression via YAP1 inhibition (Fig. 5F). The model also incorporates established components of pancreatic Notch signaling, including NEUROG3-induced *DLL1* expression,^81^ DLL1-mediated activation of Notch signaling in neighboring cells,^82^ and inhibition of *NEUROG3* and *DLL1* expression by HES1.^83,63^ Further investigation into the expression patterns of *NEUROG3*, *YAP1*, *DLL1*, *DLL3*, and *HES1* will be needed to determine how these factors are coordinated in time and space. Additionally, while unidirectional shear stress in the 2-33 mPa range upregulated multiple endocrine-associated genes in a dose-dependent manner (Fig. 4C), downregulation of *WWTR1* and YAP/TAZ target genes appeared to be dose-independent (Fig. 5B). This suggests that other mediators of shear stress effect on endocrine markers may be present.

Our findings corroborate previous research showing that physical parameters can potentially influence pancreatic progenitor cell state. *In vitro* microfluidic models have the advantage of applying well-defined fluid flow conditions and reducing confounding factors. During development, pancreatic progenitor cells experience a myriad of physical cues including shear stress, pressure, stiffness, local tissue geometry, and extracellular matrix composition. Further research is needed to elucidate how the combined effect of physical and chemical signals within the dynamic microenvironment modulates cellular and tissue behavior. Such insights could also help guide the scale-up of β-cell manufacturing for replacement therapy, as bioreactor size and agitation speed have been shown to influence the fluid shear stress exerted on cell aggregates.^84,85^ Our results suggest that hydrodynamic effects on mechanosignaling and differentiation may need to be considered when translating bench-scale differentiation protocols to large-scale bioprocessing.

### 3.6 Fluid flow influences the arrangement of pancreatic progenitor cells

In addition to changes in gene expression, we hypothesized shear stress would also modulate pancreatic progenitor cell functions. We set up a variant system and workflow compatible with live cell imaging to track cell patch boundary expansion parallel to the flow direction. Previous studies on this topic typically created cell-free zones by mechanically scratching the confluent cell layer with a plastic pipette tip,^86,87^ whereas alternative approaches such as physical exclusion method have been developed later for static culture.^88^ The barrier-based assay has been shown to result in a different cell migration speed compared to the traditional wound healing assay.^89^ In our setup (Fig. 6A), cells were initially seeded in the central area and cultured under the static condition to reach confluence. The narrow well PDMS device was then detached to remove physical exclusion permitting expansion to fibronectin-coated cell-free zones under either static or constant flow conditions (8mPa).

**Figure 6.**
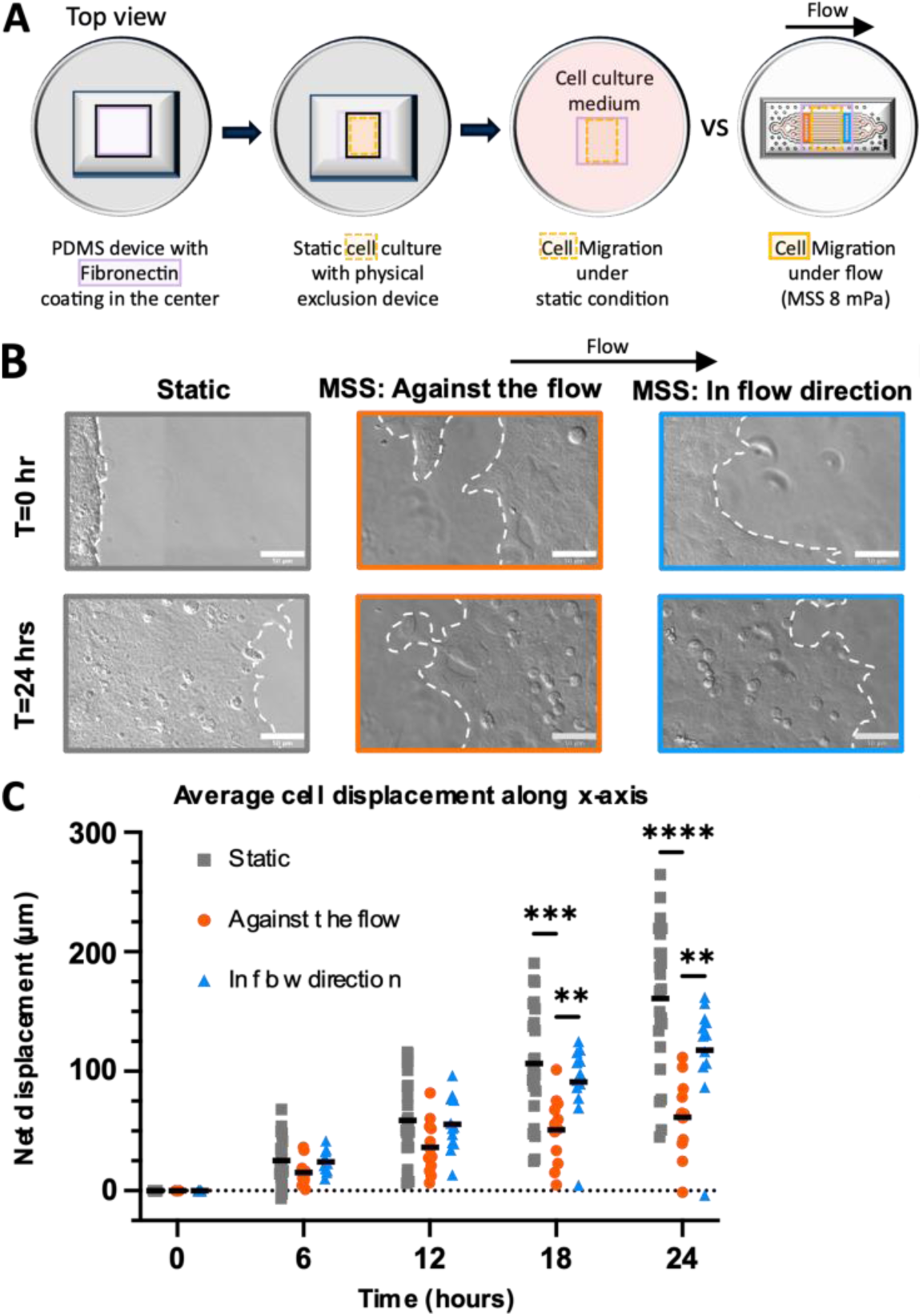
Barrier-based migration assay under shear stress. (A) Schematic of the assay workflow. (B) Representative phase-contrast images of cell patch boundaries tracked over 24 hours. Scale bar=50 mm. (C) Average net displacement distance of the cell patch boundary quantified as scattered dot plots. Static: N=24 replicates, n=3 experiments; Shear stress: N=12 replicates per flow direction, n=3 experiments. Two-way ANOVA with Tukey’s test. All conditions show significant differences across timepoints (P≤0.05). **: P≤0.01, ***: P≤0.001, ****: P≤0.0001.

We observed that cell patch boundaries expanded under all three conditions: static, against flow, and in the direction of flow (Fig. 6B). Furthermore, expansion was greater in the direction of fluid flow than against the flow (Fig. 6C). This observation demonstrated that fluid flow affects the behavior of pancreatic progenitor cells beyond changes in gene and protein expression.

## Conclusions

In this study, we investigated the effects of fluid shear stress on pancreatic progenitor cells using custom-designed microfluidic devices. By enabling mode switch between an open well device for static culture and a microchannel device for shear stress application, reversible sealing allowed the use of relatively small channel heights. This made it possible to achieve 24-hour perfusion using cost-effective and easy-to-operate syringe pumps. We then used this novel device to evaluate a novel cell type for unreported mechanosensitivity to physiologically-relevant shear stress. Pancreatic progenitor cells exhibited graded gene expression responses to the shear stress magnitudes examined (2, 8, 33 mPa), with responses to 8 mPa and 33 mPa being more similar than the response to 2 mPa. GSEA identified enrichment of cell cycle- and pancreatic endocrine-related gene sets. Notably, 24 hours of perfusion at 2 - 33 mPa induced marked upregulation of the endocrine master regulator *NEUROG3*, accompanied by downregulation of *WWTR1* gene and YAP/TAZ target genes. In addition, exposure to 8 mPa fluid shear stress reduced YAP1 protein abundance and nuclear localization, suggesting a potential role for YAP1 in fluid flow-induced *NEUROG3* upregulation. Lastly, we found pancreatic progenitor cells showed altered arrangement under fluid flow.

Overall, we demonstrated that mPa-level fluid shear stress modulates pancreatic progenitor cell signaling and function. The microfluidic system and our findings could provide a foundation for better understanding of how physical cues influence pancreatic duct development and facilitate future studies of mechanobiology in difficult-to-access tissues.

## Acknowledgement

We gratefully acknowledge funding support by DFG under Germany’s Excellence Strategy - EXC-2068 - 390729961- Cluster of Excellence Physics of Life (PoL) of TU Dresden and the Max Planck Society, as well as technical support from the Physics of Life and Center for Molecular and Cellular Bioengineering Microstructure Core Facility, PoL Microfluidics Core, PoL Microscopy Facility, and CMCB Technology Platform Flow Cytometry Core Facility. We also acknowledge technical support with initial cell expansion prior to cryopreservation from Marit Leuschner and thank Coline Schewin for insights regarding ductal diameters and flow rates inside mouse embryonic pancreas. Figure 1D (tm8fmit) and 5F (c7kfumv) schematics were created in BioRender (licenses: Yu, X. 2026). ChatGPT (GPT-5.5) was used to assist in revising the manuscript for clarity and generating code for transcriptomic analyses. All AI-generated text and code were iteratively reviewed, revised, and verified by the authors, who made all methodological and analytical decisions.

## Data availability

The data supporting this article are included as part of the electronic supplementary information.

