## Supplemental Figures for "Pancreatic progenitor cells respond to laminar shear stress with transcriptomic changes, accompanied by reduced YAP1 activity"

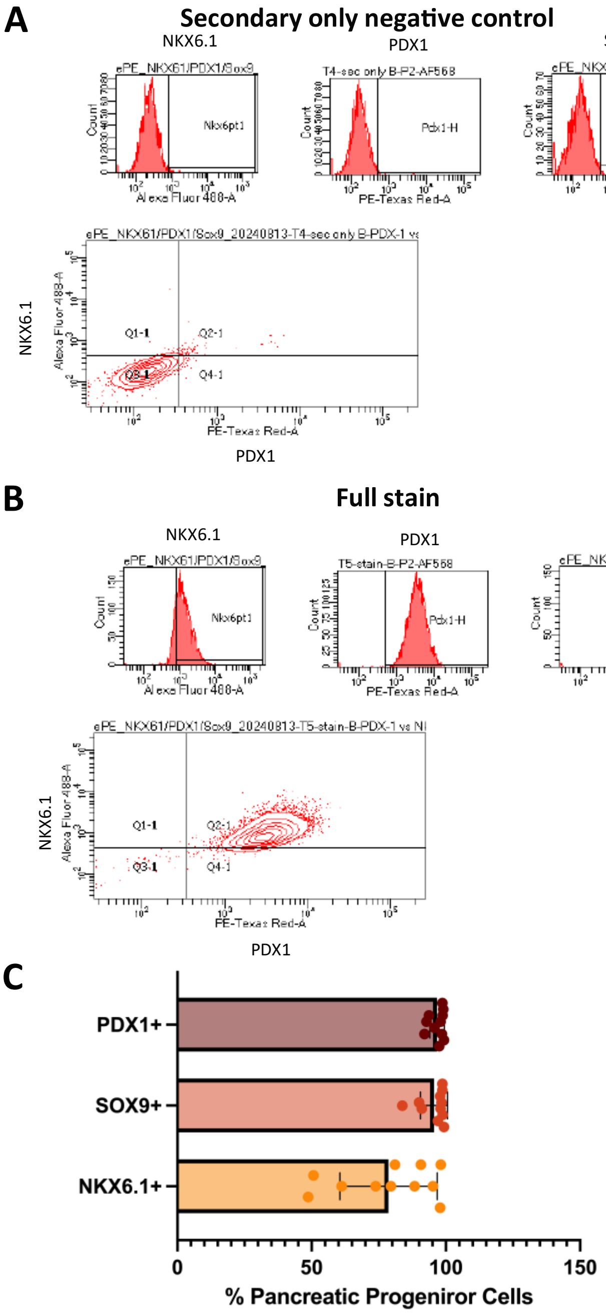

Figure S1. Flow cytometry analysis of progenitor cell markers expressed in ePE cells. (A) Representative measurement of a negative control stained with secondary antibodies only. (B) Representative measurement of a sample stained with primary and secondary antibodies. (C) Average percentage of $PDX1^{+}$, $SOX9^{+}$, $NKX6.1^{+}$ ePE cells across n=11 experiments.

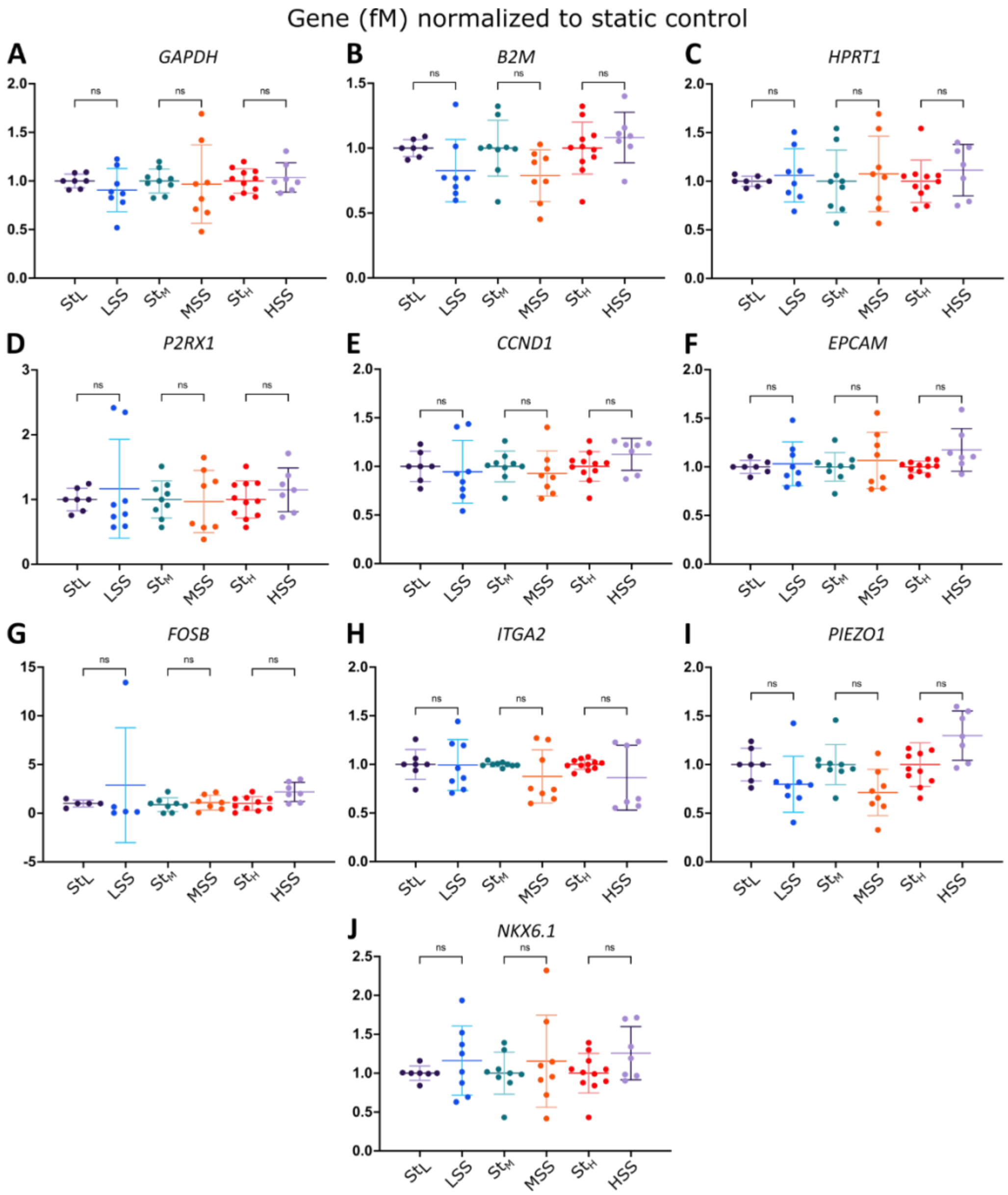

Figure S2. qPCR measurement and analysis for (A-C) housekeeping genes, (D-I) remaining of the 13 putative shear-responsive candidate genes, and (J) a pancreatic progenitor cell marker. N≥7 replicates per condition, n=4 experiments per SS-vs-Static comparison. Brown-Forsythe and Welch ANOVA with Dunnett T3 post hoc test. ns: P>0.05.

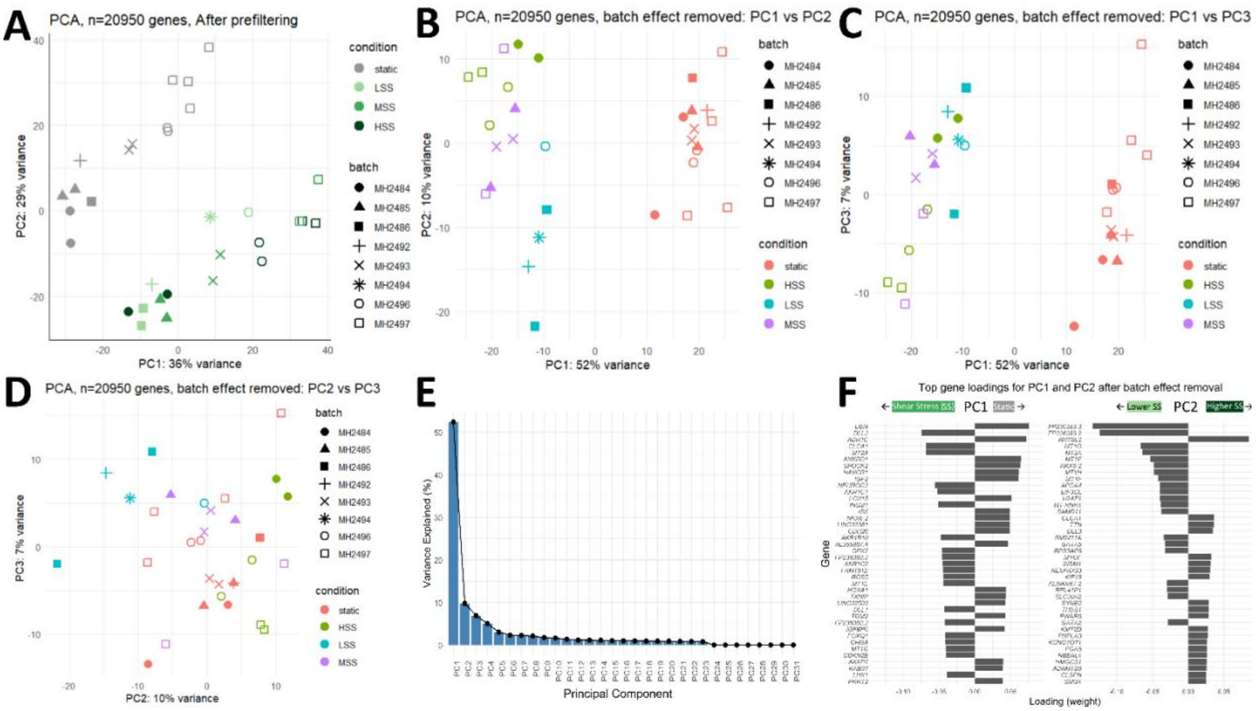

G

| | Genes | LSS vs static padj | LSS vs static log2FC_shrunk | MSS vs static padj | MSS vs static log2FC_shrunk | HSS vs static padj | HSS vs static log2FC_shrunk |
|---|---|---|---|---|---|---|---|
| **Responsive** | *EGR1* | 1.26E-08 | 1.11 | 7.41E-25 | 1.40 | 6.14E-44 | 1.83 |
| | *KRT19* | 6.20E-03 | 0.34 | 4.51E-03 | 0.26 | 4.17E-05 | 0.36 |
| | *FOS* | 6.55E-09 | 0.94 | 3.75E-24 | 1.15 | 2.43E-39 | 1.46 |
| | *SDC1* | 9.25E-02 | 0.23 | 3.38E-04 | 0.35 | 5.89E-04 | 0.34 |
| | *CXCL12* | 2.97E-07 | 0.68 | 2.23E-18 | 0.83 | 1.67E-28 | 1.03 |
| | *ACTB* | 2.94E-03 | 0.31 | 2.83E-06 | 0.35 | 6.83E-07 | 0.36 |
| | *CD44* | 9.36E-03 | 1.05 | 1.84E-03 | 1.27 | 3.90E-04 | 1.43 |
| | *TFPI* | 2.63E-06 | -0.49 | 4.78E-17 | -0.62 | 7.22E-31 | -0.83 |
| **Non-responsive** | *CCND1* | 9.61E-02 | -0.16 | 6.61E-01 | -0.04 | 7.00E-02 | -0.13 |
| | *EPCAM* | 7.70E-01 | 0.04 | 9.24E-01 | 0.01 | 9.90E-01 | 0.00 |
| | *FOSB* | 9.79E-01 | 0.01 | 8.86E-01 | 0.03 | 8.86E-01 | -0.03 |
| | *ITGA2* | 1.76E-01 | -0.21 | 6.40E-03 | -0.31 | 4.63E-05 | -0.45 |
| | *P2RX1* | 9.49E-01 | 0.01 | 2.19E-01 | 0.26 | 1.78E-01 | 0.31 |
| | *PIEZO1* | 2.26E-01 | -0.18 | 2.61E-02 | -0.27 | 2.34E-01 | -0.15 |
| **Progenitor marker** | *NKX6-1* | 1.26E-01 | 0.26 | 1.17E-02 | 0.33 | 8.18E-01 | 0.04 |
| **Housekeeping genes** | *GAPDH* | 5.78E-01 | 0.06 | 7.37E-01 | 0.03 | 3.21E-01 | 0.08 |
| | *B2M* | 8.97E-01 | 0.02 | 1.46E-01 | -0.10 | 3.26E-03 | -0.18 |
| | *HPRT1* | 1.16E-01 | 0.20 | 7.96E-02 | 0.16 | 1.83E-01 | 0.13 |

Figure S3. Transcriptomic analysis of bulk RNA-seq data. (A) Principal component analysis (PCA) of 31 samples based on 20950 expressed genes before batch effect correction. (B-D) PCA plots after batch effect correction. (E) Scree plot showing the variance explained by each principal component after batch effect correction. (F) Top loading genes contributing to PC1 and PC2. (G) Bulk RNA-seq expression of putative shear-responsive gene candidates previously measured by qPCR. Red: log2FC shrunk > 0; Blue: log2FC shrunk < 0; Purple: $P_{adj}$<0.05.

A

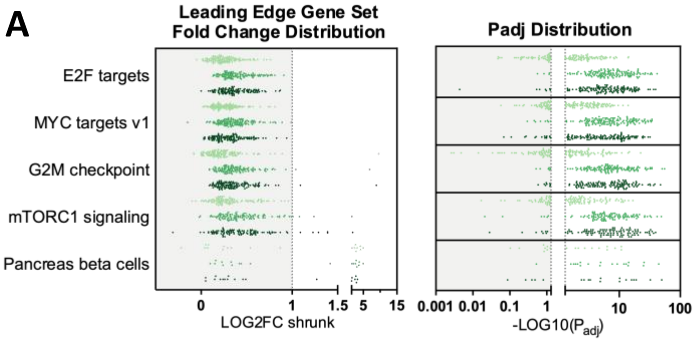

B

| | log2FC shrunk | | | -log10(Padj) | | |
|---|---|---|---|---|---|---|
| Genes | L vs St | M vs St | H vs St | L vs St | M vs St | H vs St |
| *CCNA1* | 0.7803 | 1.2134 | 0.7692 | 0.80 | 5.80 | 2.90 |
| *CDKN2B* | 1.7595 | 2.2532 | 2.6565 | 0.17 | 21.47 | 30.68 |
| *GADD45G* | 0.2542 | 1.3989 | 2.0078 | 1.18 | 6.80 | 13.60 |
| *SFN* | 0.0241 | 0.9303 | 1.1315 | 1.50 | 2.67 | 3.65 |

log2FC shrunk: 2.66 – 0
-log10(Padj): ≥1.3; (0, 1.3)

C

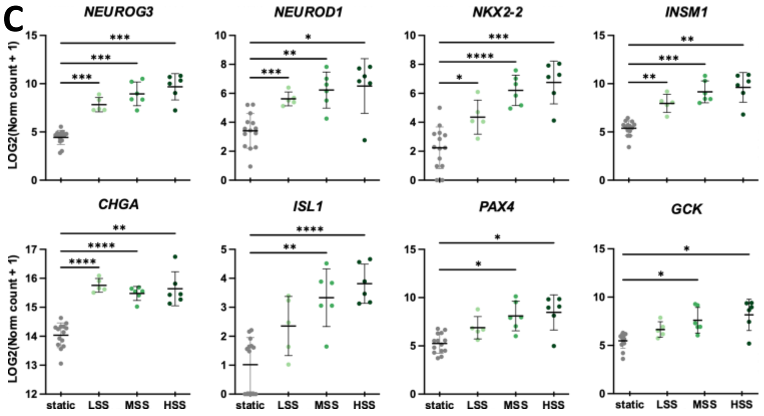

D

| | | | | Intersection Size | | | | -log10(Padj) | | |
|---|---|---|---|---|---|---|---|---|---|---|
| Source | Term name | Term id | Term size | L | M | H | Union | L | M | H |
| GO:MF | signaling receptor activity | GO:0038023 | 1009 | 64 | 122 | 124 | 161 | 6.45 | 16.91 | 15.00 |
| | molecular transducer activity | GO:0060089 | 1009 | 64 | 122 | 124 | 161 | 6.45 | 16.91 | 15.00 |
| | transmembrane signaling receptor activity | GO:0004888 | 546 | 33 | 70 | 71 | 90 | 0.72 | 8.84 | 7.71 |
| | receptor ligand activity | GO:0048018 | 298 | 26 | 43 | 47 | 60 | 2.64 | 5.41 | 6.55 |
| | signaling receptor activator activity | GO:0030546 | 304 | 26 | 43 | 47 | 60 | 2.47 | 5.13 | 6.24 |
| | signaling receptor regulator activity | GO:0030545 | 325 | 26 | 43 | 47 | 60 | 1.90 | 4.20 | 5.21 |
| | gated channel activity | GO:0022836 | 222 | 22 | 32 | 34 | 43 | 2.57 | 2.96 | 3.27 |
| | hormone activity | GO:0005179 | 72 | 13 | 17 | 17 | 20 | 2.86 | 2.77 | 2.30 |
| GO:CC | extracellular matrix | GO:0031012 | 422 | 19 | 52 | 62 | 74 | 0.00 | 4.96 | 8.68 |
| | external encapsulating structure | GO:0030312 | 423 | 20 | 52 | 62 | 75 | 0.00 | 4.92 | 8.63 |
| | collagen-containing extracellular matrix | GO:0062023 | 303 | 12 | 38 | 44 | 54 | 0.00 | 2.71 | 4.76 |
| GO:BP | system process | GO:0003008 | 1441 | 65 | 128 | 136 | 172 | 1.08 | 7.33 | 7.45 |
| | G protein-coupled receptor signaling pathway | GO:0007186 | 558 | 30 | 61 | 66 | 82 | 0.00 | 4.37 | 5.09 |
| | cell motility | GO:0048870 | 1437 | 56 | 113 | 125 | 165 | 0.00 | 2.75 | 4.10 |
| | metal ion transport | GO:0030001 | 660 | 39 | 64 | 69 | 90 | 1.57 | 2.73 | 3.22 |
| | cell fate determination | GO:0001709 | 30 | 8 | 11 | 12 | 13 | 1.59 | 2.51 | 3.24 |

Intersection Size: 172 – 8
-log10($P_{adj}$): 10 – 1.3

Figure S4. ≥Functional enrichment analyses. (A) LOG2FC shrunk and -LOG10($P_{adj}$) of leading-edge genes in the top 5 enriched hallmark gene sets under shear stress conditions. (B) KEGG cell cycle genes that met DEG criteria. (C) normalized expression of endocrine-related genes (no batch effect correction). (D) Number of DEGs overlapping with top overrepresented gene ontologies. Brown-Forsythe and Welch ANOVA with Games-Howell post hoc test. *: P≤0.05, **: P≤0.01, ***: P≤0.001, ****: P≤0.0001.

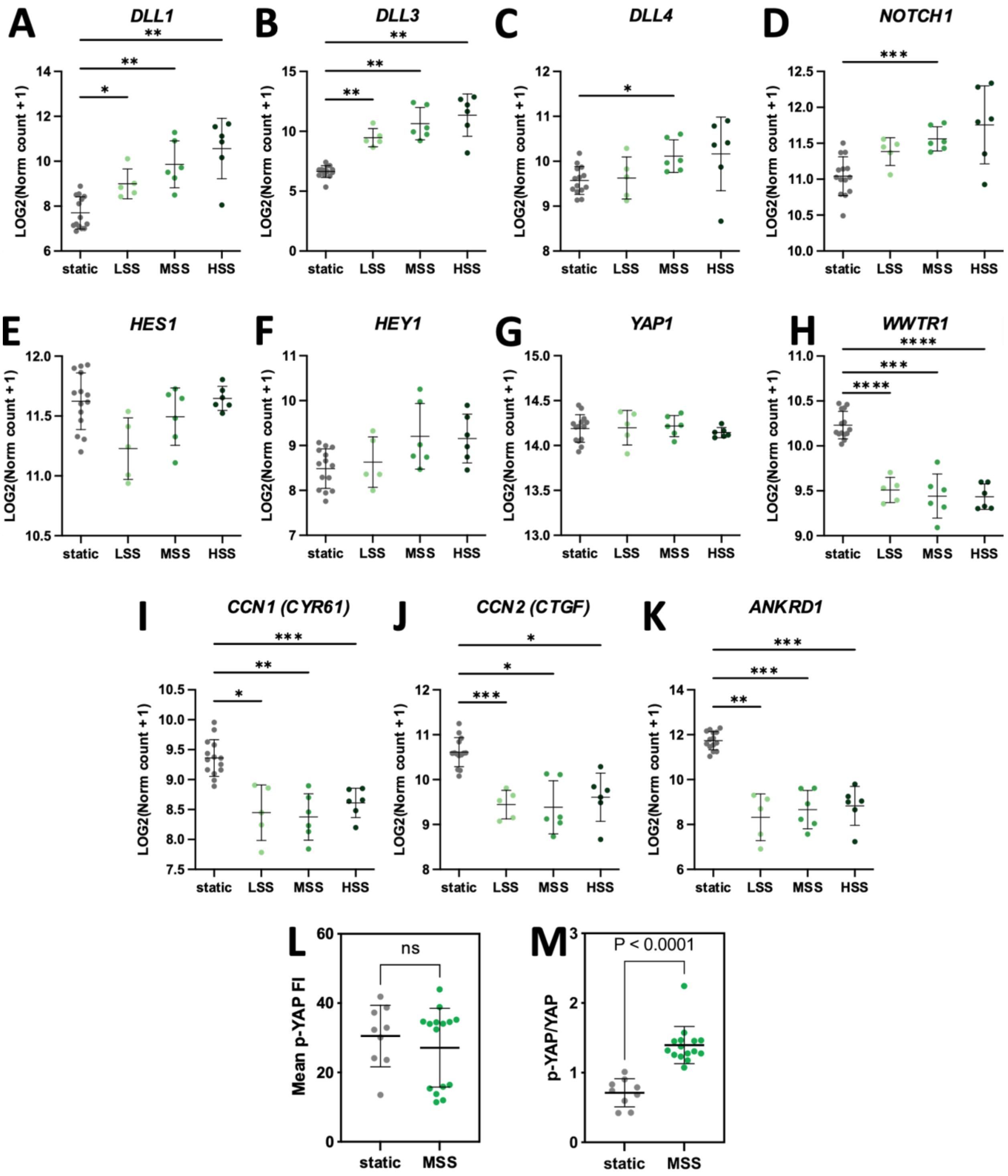

Figure S5. Evaluation of NOTCH and YAP signaling. (A-K) Normalized expression from RNA-seq data without batch effect correction. Brown-Forsythe and Welch ANOVA with Games-Howell post hoc test. *: P≤0.05, **: P≤0.01, ***: P≤0.001, ****: P≤0.0001. (L-M) Scatterplots of mean phosphorylated YAP1 (Ser127) fluorescence intensity (FI) and its ratio to YAP1 FI. Welch's t-test. ns: not significant.
