## Supplemental Methods for "Pancreatic progenitor cells respond to laminar shear stress with transcriptomic changes, accompanied by reduced YAP1 activity"

**Supplemental method: Calculation of oxygen content**

Oxygen content was calculated using steady-state mass balance incorporating convective delivery, transport through the PDMS layer, and cellular consumption. Assuming that the medium at the channel inlet and PDMS exterior were at the same oxygen equilibrium, the outlet oxygen concentration was calculated using the following equations:

convective output = convective input + transport through PDMS − cellular uptake

$$Q(C + \frac{dC}{dx}dx) = QC + k_c w dx(C_{ext} - C) - r dx \quad (1)$$

C = C(x) = cross-section-average dissolved oxygen species,

x = position along the cell path;

$C_{ext}$ = equivalent dissolved oxygen concentration corresponding to the external incubator atmosphere;

w = channel width;

Q = flow rate;

$k_c$ = effective oxygen-transfer coefficient based on dissolved concentration;

r = cellular consumption per channel length.

$$Q\frac{dC}{dx} = k_c w(C_{ext} - C) - r$$

This equation is a variant form of the equation 4 from Dharia et al (2018).[1] We also added a cellular consumption term here.

$$\frac{dC}{dx} + \frac{k_c w}{Q}C = \frac{k_c w}{Q}C_{ext} - \frac{r}{Q}$$

Define $A_{PDMS} = wL$ with $L$ being the length of the channel, and $K = k_c A_{PDMS}$,

$$\frac{dC}{dx} + \frac{K}{QL}C = \frac{K}{QL}C_{ext} - \frac{r_{channel}}{QL} \quad (2)$$

where $r_{channel} = rL$

When $\frac{dC}{dx}$=0, $C_{constant}$= $C_{ext} - \frac{r_{channel}}{K}$

At inlet, we have the boundary condition C(0) = $C_{in}$

$$C(x) = C_{constant} + (C_{in} - C_{constant})e^{-\frac{Kx}{QL}}$$

$$C(x) = C_{ext} - \frac{r_{channel}}{K} + (C_{in} - C_{ext} + \frac{r_{channel}}{K})e^{-\frac{Kx}{QL}}$$

At outlet, $C_{ext}$=$C_{in}$

$$C_{out} = C_{in} - \frac{r_{channel}}{K}[1 - e^{-\frac{K}{Q}}] \quad (3)$$

The effective oxygen-transfer conductance was estimated as:

$$K = \frac{A_{PDMS}}{\frac{S_{media}h_{PDMS}}{P_{PDMS}} + \frac{h_{channel}}{D_{media}}} \quad (4)$$

| Parameter | Description | Value | source |
|---|---|---|---|
| $C_{in}$ | Inlet oxygen concentration | 0.17075 mol $m^{-3}$, corresponding to approximately 17% $O_2$ | Ochs et al (2014) supplement[2] |
| $Q$ | Volumetric flow rate | 0.17 µL $min^{-1}$ at LSS<br>0.67 µL $min^{-1}$ at MSS<br>2.75 µL $min^{-1}$ at HSS | Set in this study to achieve desired shear stress |
| $r_{channel}$ | Total cellular oxygen uptake rate per channel | 0.64 pmol $s^{-1}$ | Calculated from 0.016pmol/s per 1500 cells[3] assuming we have 60K cells per channel |
| $A_{PDMS}$ | Area of PDMS-medium interface (per channel) | 0.08$cm^2$ | Set in this study |
| $h_{PDMS}$ | PDMS thickness | 5mm | Set in this study |
| $P_{PDMS}$ | Oxygen permeability in PDMS | 732.7 barr or 2.45x$10^{-13}$ mol/m/s/Pa | Sønstevold et al (2023)[4] |
| $h_{channel}$ | Channel height | 100 µm | Set in this study |
| $D_{media}$ | Diffusion coefficient of oxygen in media | 3.37 x $10^{-5}$ $cm^2/s$ | Petterson et al (2005)[5] |
| $S_{media}$ | Oxygen solubility in media | 9.72 x $10^{-6}$ mol/$m^3$/Pa | Komlódi et al (2021)[6] |

$$K = 3.508 \times 10^{-11}\ m^3/s = 2.1\mu L/min$$

Oxygen % can then be calculated using $\%Oxygen = \frac{C_{out}/S_{media}}{P_{atm}}$, where $P_{atm}$ = 101325 Pa.

| Shear stress | $C_{out}$ | $\%Oxygen$ |
|---|---|---|
| LSS | 0.15251 $mol/m^3$ | 15.5% |
| MSS | 0.15330 $mol/m^3$ | 15.6% |
| HSS | 0.16099 $mol/m^3$ | 16.3% |

The calculated outlet oxygen levels were approximately 15.5%, 15.6%, and 16.3% under LSS, MSS, and HSS, respectively.

**Supplemental method references:**
